# Characterizing psilocybin as an antidepressant for adolescence in male and female rats

**DOI:** 10.1101/2024.12.20.629571

**Authors:** Rubén García-Cabrerizo, Itziar Beruete-Fresnillo, M. Julia García-Fuster

**Author notes:** **Correspondence:** and University of the Balearic Islands.

## Abstract

Adolescent depression is a significant public health concern, yet treatment options remain limited, particularly due to age- and sex-related differences in antidepressant efficacy. This study explored the rapid and long-lasting antidepressant-like potential of psilocybin in adolescent Sprague-Dawley rats, examining acute and repeated oral dosing effects while incorporating sex as a biological variable. An acute administration of psilocybin produced rapid antidepressant-like effects 30 minutes post-treatment in both male and female rats, demonstrated by reduced immobility and increased escape-related behaviour in the forced swim test. However, repeated daily administrations over 7 days revealed notable sex differences. In males, the antidepressant-like effects were sustained, at least, for up to 15 days post-treatment at both tested doses. In contrast, in females, the effects were dose-dependent and less enduring, persisting only up to 8 days at the highest dose tested. To the best of our knowledge, these results are the first ones to underscore psilocybin’s potential as a fast-acting and long-lasting antidepressant during adolescence, a developmental stage marked by high vulnerability to depression and reduced response to conventional treatments, while also emphasizing the importance of tailoring therapeutic approaches to individual biological factors such as sex.

## 1. Introduction

Age and sex play a key role in the diagnosis and treatment of major depression, as they significantly influence the disorder’s prevalence. Major depressive disorder is a widespread mental health concern among adolescents, affecting about 5-6% of this group (Costello et al., 2006), with rates higher in females during adolescence and continuing into adulthood (Kessler, 2003; LeGates et al., 2019; Marcus et al., 2005). Recently, adolescent depression has increased due to factors such as academic pressure, high expectations, peer influence, and the mental health impact of the COVID-19 pandemic (Moreno et al., 2020; Pratibha and Virender Kumar, 2020). Despite the growing vulnerability of adolescents to depression, available treatment options remain limited, with fluoxetine and escitalopram as main options in combination with psychotherapy (Boaden et al., 2020). The limited availability of safe pharmacological treatments for adolescents may be linked to age-related differences in the underlying biology of depression, as the adolescent brain is still in a developmental phase, which could contribute to the generally weaker response to antidepressants compared to adults (Bylund and Reed, 2007). Moreover, given the increased risk of suicidal behaviour among adolescent with depression, there is an urgent demand to identify new, fast-acting antidepressants tailored for this vulnerable population (Ledesma-Corvi et al., 2024).

The tryptamine alkaloid psilocybin (4-phosphoryloxy-N,N-dimethyltryptamine) is a psychoactive substance known for its ability to alter perception, mood, and cognitive processes (Nichols, 2004). Some of these effects are thought to be mediated by its binding to the 5-HT_2A_ serotonin receptor, and previous research suggests that psilocybin use does not typically lead to substance dependence (Kelmendi et al., 2022). In recent years, several research efforts have focused on exploring the therapeutic potential of micro-dosed psilocybin for a variety of health conditions, such as substance use disorders, alleviating cancer-related mental health issues, and as a promising treatment for post-traumatic stress disorder, anxiety and depression (Lea et al., 2020; Rootman et al., 2021). In support of these ongoing studies, the FDA has granted a “Breakthrough Therapy” designation to two psilocybin formulations being evaluated for their safety and effectiveness as medical treatments for adult depression (Nutt et al., 2020). However, there remain significant questions about the potential effectiveness of psilocybin on the developing adolescent brain and the impact of its use as antidepressant, highlighting the critical need for further pre-clinical assessments.

Clinical trials suggest that psilocybin, especially when combined with psychotherapy, holds strong potential for treating depression, including major depressive disorder and treatment-resistant depression (Carhart-Harris et al., 2021; Goodwin et al., 2023; Gukasyan et al., 2022; Raison et al., 2023; Rosenblat et al., 2024; Sloshower et al., 2023; von Rotz et al., 2023). It is considered safer than other psychedelics, with low physiological toxicity, minimal risk of addiction, and generally mild side effects such as headaches and nausea (Bogenschutz et al., 2015; Lowe et al., 2021). Research has shown that both single and multiple high doses can significantly alleviate depressive symptoms, and in some cases, it may be more effective than conventional antidepressants (Carhart-Harris et al., 2021). Although there are challenges in keeping participants blinded due to psilocybin’s distinct effects, its promising safety and efficacy make it a candidate worth reconsidering for broader clinical use and regulatory adjustments. Despite this evidence, its effectiveness during adolescence has been largely unexplored. Therefore, the present study aims to investigate the antidepressant potential of psilocybin in adolescents, incorporating sex as a biological variable. In particular, the research will assess the immediate and long-term antidepressant-like effects of various doses of psilocybin in both male and female adolescent rats.

## 2. Materials and methods

### Animals and drug treatment

A total of 60 adolescent male and female Sprague-Dawley rats (30 males, 30 females) were bred in the animal facility at the University of the Balearic Islands. All procedures were performed following the ARRIVE guidelines (Percie du Sert et al., 2020), the EU Directive 2010/63/EU of the European Parliament and of the Council, after approval by the Local Committee (University of the Balearic Islands; CEEA: 235-04-24) and the regional Government (Conselleria de Medi Ambient, Agricultura i Pesca, Direcció General Agricultura i Ramaderia, Govern de les Illes Balears AEXP: SSBA 12/2024). All efforts were made to minimize the number of rats used and their suffering. The procedures were performed during the light period (between 09:00 h and 13:00 h). At post-natal day (PND) 21, animals were housed (22 °C, 70% humidity, 12:12 h light /dark cycle, lights on at 08:00 a.m. and of at 20:00 p.m.) in standard cages in groups of 2-4 animals with *ad libitum* food and tap water. Animals were handled for 1 day (PND 33) before being treated with a daily dose of psilocybin (0.3 or 1 mg/kg prepared in saline and administered by oral gavage; Lipomed AG, Arlesheim, Switzerland) or saline (0.9% NaCl, 2 ml/kg by oral gavage) for 7 consecutive days (PND 35-41, see Figure 1A). The license to conduct research with psilocybin was given by AEMPS (register number: 00084411837). These doses of psilocybin used were based on prior studies showing behavioural and neurochemical effects of diverse psychedelics in rodents (Erkizia-Santamaría et al., 2022; Inserra et al., 2023; Takaba et al., 2024; Zhao et al., 2024). Animals were weighted daily thought treatment administration.

**Figure 1.**
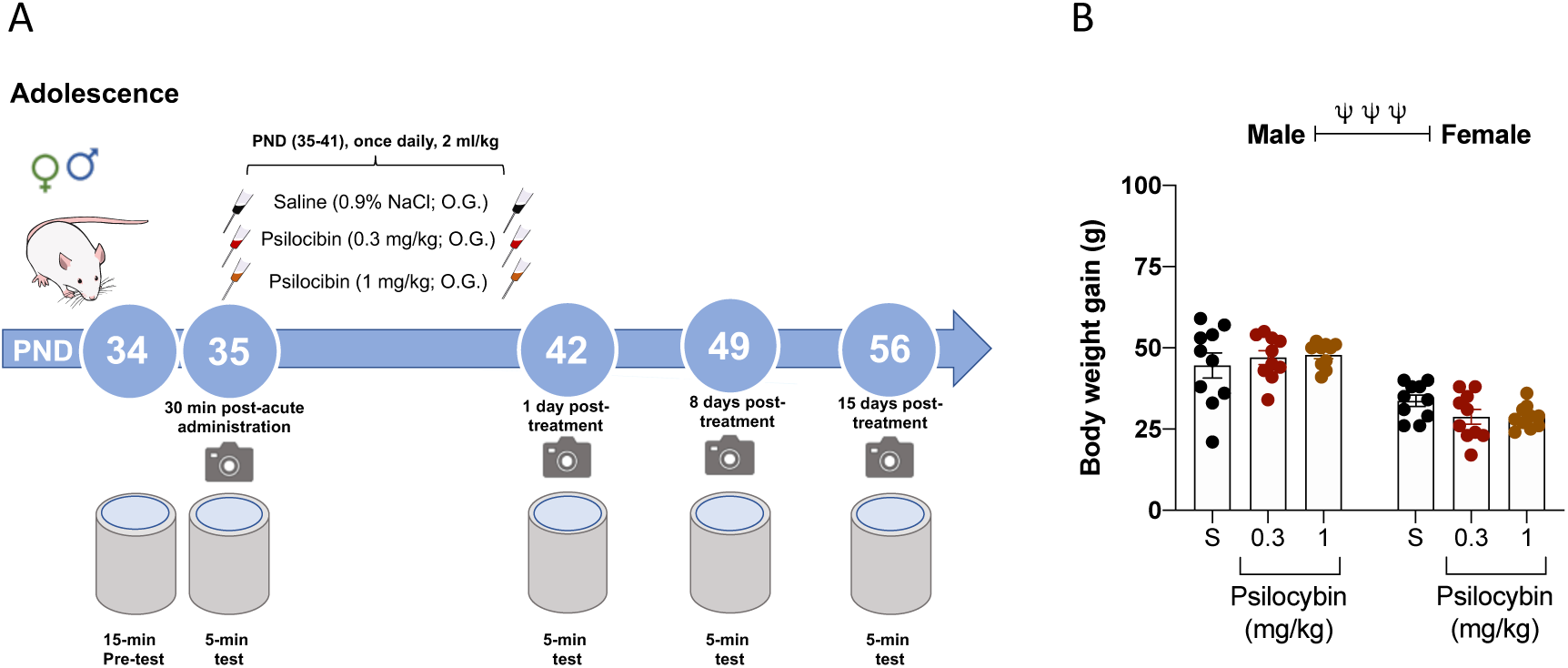
Experimental design. A) Behavioural procedures in adolescent male and female rats after an acute (1 dose of 0.3 and 1 mg/kg, oral gavage, O.G.) or a repeated (7 doses of 0.3 and 1 mg/kg, O.G., 1 dose per day) treatment of psilocybin. Changes were evaluated 30-min (post-natal day, PND, 35) or 1-, 8- and 15-days post-treatment (PND 42, 49 and 56) in the forced-swim test (FST). B) Changes in body weight gain during treatment with psilocybin in adolescent male and female rats. Columns represent mean ± SEM of body weight gain (g). Individual symbols are shown for each rat. Two-way ANOVA (independent variables: Sex and Treatment) ^ΨΨΨ^*p*<0.001 when comparing male vs female rats (Effect of Sex).

### Forced-swim test

To assess the potential antidepressant-like efficacy of psilocybin in adolescent male and female rats we relied on the forced-swim test (FST), a validated behavioural tool in naïve rodents to ascertain antidepressant-like responses under stress conditions (Slattery and Cryan, 2012). Each rat was individually placed in a water tank (41 cm high x 32 cm diameter, 25 cm depth) at 25 ± 1 °C during 15 min (pre-test session, PND 34) followed, 24 h later, by a 5 min test session 30 minutes after the first psilocybin administration at PND 35 that was videotaped (García-Cabrerizo et al., 2015; García-Cabrerizo and García-Fuster, 2019). Videos were blindly analysed to determine individual levels of immobility (defined as the lack of movement except that which is necessary to keep the rat’s nose above the water level), climbing or swimming for each rat (Behavioral Tracker software, CA, USA). Then, the antidepressant-like effect induced by repeated psilocybin was evaluated at different time-points after treatment (1, 8 and 15 days) by re-exposing rats to 5 min sessions in the FST (see Figure 1A). Similarly, prior studies from our group repeatedly exposed rats to the FST across time, demonstrating that the possible effects of learning due to repetition can be controlled if the same conditions are applied to all rats, and that this set-up allows for reliable measurements of the progression of this particular behavioural response (Bis-Humbert, García-Cabrerizo and M Julia García-Fuster, 2021; García-Cabrerizo et al., 2020; García-Cabrerizo and García-Fuster, 2019; Jiménez-Romero et al., 2020).

### Statistical analysis

Data was analysed with GraphPad Prism, Version 10 (GraphPad Software, Inc., California, USA). Results are expressed as mean values ± standard error of the mean (SEM) and individual symbols are shown for each rat. Each set of data, including changes in body weight gain, time spent immobile, climbing, or swimming in the FST, was evaluated using two-way analyses of variance (ANOVAs) with independent variables such as Sex (male vs. female) and Treatment (saline vs. psilocybin), followed by Tukey’s multiple comparisons test when appropriate. The level of significance was set at *p* ≤ 0.05.

## 3. Results

### Body weight

Body weight gain (g) was monitored during treatment administration (Figure 1B). A two-way ANOVA did not detect an effect of Treatment (F_2,54_=0.16, *p*=0.852), however it detected a significant effect of Sex (F_1,54_=76.49, *p*<0.0001); demonstrating a lower body weight gain in adolescent female rats as expected.

### Acute antidepressant-like effects induced by psilocybin in males and female rats

The immediate effects of psilocybin were assessed in the FST 30 min after a single oral dose (Figure 2A-C). A two-way ANOVA revealed a significant impact of Treatment (F_2,54_=18.8, *p*<0.0001). Specifically, both doses of psilocybin significantly decreased the time spent immobile in the FST when compared to saline-treated rats, in both male (-48 ± 14 s, S vs. 0.3 mg/kg, \*\**p*=0.0028; -69 ± 14 s, S vs. 1 mg/kg, \*\*\**p*<0.001) and females (-37 ± 14 s, S vs. 0.3 mg/kg, \**p*=0.0275; -46 ± 14 s, S vs. 1 mg/kg, \*\**p*=0.0045) (Figure 2A). These decreases in immobility were paired with increases in active escape behaviours, such as climbing and swimming, indicating that psilocybin may have a rapid antidepressant-like effect (Figure 2B, C).

**Figure 2.**
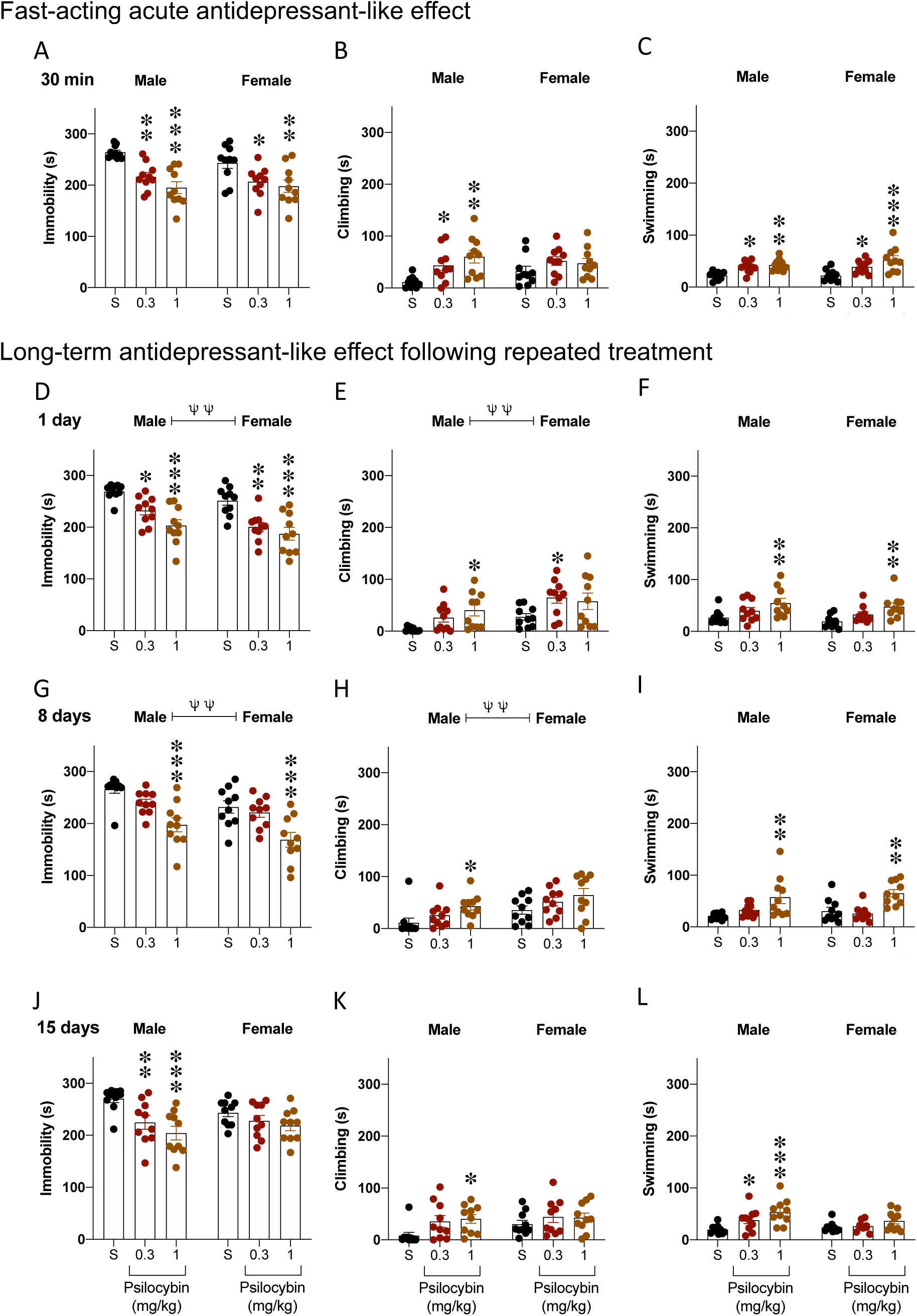
Evaluating the potential antidepressant-like effects of different doses of psilocybin in adolescent male and female rats. A-C) Fast-acting acute antidepressant-like effects of psilocybin (1 single dose of 0.3 and 1 mg/kg, oral gavage, O.G; post-natal day, PND, 35) as measured in the forced swim-test (FST) 30 min post-treatment. D-L) Long-term antidepressant-like effect following repeated treatment with psilocybin (0.3 and 1 mg/kg, O.G; 7 days, PND 35-41) as evaluated in the FST 1-, 8- and 15-days post-treatment (PND 42, 49 and 56). Columns represent mean ± SEM of time spent (s) in each behaviour (immobility, climbing and swimming). Individual symbols are shown for each rat. Two-way ANOVA (independent variables: Sex and Treatment) followed by Tukey’s multiple comparisons test; ^ΨΨ^*p*<0.01 when comparing male vs female rats (Effect of Sex); \**p* < 0.05, \*\**p* < 0.01, \*\*\**p* < 0.001 when comparing psilocybin-treated rats (0.3 and 1 mg/kg) vs. saline-treated rats.

### Sex differences in the antidepressant-like effects induced by repeated psilocybin exposure during adolescence

To assess the antidepressant-like effects of repeated psilocybin treatment at different time points post-intervention, we measured immobility and escape behaviours in the FST. In particular, 1 day after repeated treatment, results showed a reduction in immobility time in both male and female rats (Figure 2D). A two-way ANOVA revealed significant effects of Treatment (F_2,54_=24.8, *p*<0.0001) and Sex (F_1,54_=8.1, *p*=0.006), with both doses of psilocybin significantly reducing immobility time in males (-37 ± 13 s, S vs. 0.3 mg/kg, \**p*=0.0218; -66 ± 13 s, S vs. 1 mg/kg; \*\*\**p*<0.001) and females (-51 ± 13 s, S vs. 0.3 mg/kg, \*\**p*=0.001; -64 ± 13 s, S vs. 1 mg/kg, \*\*\**p*<0.001). In this line, psilocybin increased active escape behaviours for both sexes (see Figure 2E, F). When evaluated 8 days post-treatment (Figure 2G), a two-way ANOVA again detected significant effects of Treatment (F_2,54_=19.1, *p*<0.0001) and Sex (F_1,54_=9.3; *p*=0.004). At this time point, only the highest dose of psilocybin significantly reduced immobility time in males (-69 ± 16 s, S vs. 1 mg/kg, \*\*\**p*<0.001) and females (-63 ± 16 s, S vs. 1 mg/kg, \*\*\**p*<0.001), while also increasing escape behaviours in both sexes (Figures 2H, I). Finally, at 15 days post-administration, a two-way ANOVA demonstrated a long-lasting antidepressant-like effect of Treatment (F_2,54_=9.7, *p*=0.0003) (Figure 2J). However, *post-hoc* analysis indicated a significant reduction in immobility only in male rats, with both doses of psilocybin showing significant effects compared to saline (-45 ± 15 s, S vs. 0.3 mg/kg, \*\**p*=0.0096; 66 ± 15 s, S vs. 1 mg/kg, \*\*\**p*<0.001), whereas no significant changes were observed in females. These findings align with the effects seen in active escape behaviours, where psilocybin increased these behaviours only in males (Figure 2K, L), suggesting a faster gradual loss of psilocybin’s antidepressant-like efficacy in female rats over time.

## 4. Discussion

This study demonstrated fast- and sex-specific long-lasting antidepressant-like effects induced by oral psilocybin in adolescent Sprague-Dawley rats. Psilocybin produced a rapid antidepressant-like response 30 min after administration in both male and female adolescent rats. However, sex differences were observed in the long-term antidepressant effectiveness following repeated administration. Specifically, a sustained antidepressant-like effect was observed in male rats exposed to repeated psilocybin, still lasting up to 15 days after treatment. In contrast, this antidepressant-like effect persisted for only up to 8 days after treatment in adolescent female rats, and only with the highest psilocybin dose tested. These findings underscore the critical need to consider sex as key factor in developing effective, personalized therapeutic strategies for adolescent depression, emphasizing the need for sex-tailored clinical trials and therapeutics.

Acute oral psilocybin reduced FST immobility and increased climbing and swimming behaviours within 30 min, demonstrating rapid antidepressant-like effects in both male and female adolescent rats. To date, there is a notable lack of studies specifically investigating the acute effects of a single psilocybin administration during adolescence, particularly at this early time point. Antidepressant-like effects have been reported in adult male mice 24 hours after a single intraperitoneal administration of psilocin, the active metabolite of psilocybin, assessed using the FST (Takaba et al., 2024). Similarly, sub-acute doses of a different tryptamine psychedelic, N,N-dimethyltryptamine (DMT), the principle hallucinogenic component of ayahuasca, reduced immobility in the FST 1 hour after the last administration (Cameron and Olson, 2018). In contrast, 5-methoxy-N,N-dimethyltryptamine (5-MeO-DMT), did not produce immediate antidepressant-like effects 30 min after administration in the FST, but showed delayed benefits 24 hours later in adult male mice (Cameron et al., 2023). Similar results were observed in chronically stressed male mice, were a single injection of psilocybin reversed anhedonia 24 hours post-administration but had no effect in unstressed mice of either sex (Hesselgrave et al., 2021). In contrast, a more recent study reported a rapid antidepressant-like effect in the FST following a single intraperitoneal administration of various doses of psilocybin in naïve adult male mice (Zhao et al., 2024). Consistent with our findings, substances like ketamine have demonstrated rapid antidepressant-like effects in the FST 30 minutes after administration, both in aged male rats (Hernández-Hernández et al., 2024) and in adolescent rats (Jornet-Plaza et al., 2024; Ledesma-Corvi et al., 2022). However, in adolescents, these effects were influenced by prior stress exposure, with drops in efficacy and the need for higher drug doses when males and females are subjected to early-life stress (Jornet-Plaza et al., 2024; Ledesma-Corvi et al., 2022). Therefore, and to the best of our knowledge, this is the first study to report an antidepressant-like response following an oral administration of psylocibin in adolescent rats of both sexes.

Findings obtained in the present work revealed clear sex differences in the long-term antidepressant-like efficacy following the repeated administration of psilocybin. In adolescent male rats, both doses tested significantly reduced immobility time, with effects still lasting up to 15 days post-treatment. At the highest dose, this reduction was also linked to an increase in escape behaviours, indicating a prolonged antidepressant-like response. In contrast, the effects in adolescent female rats were less enduring. While both doses produced a clear impact 1 day after treatment, the effect of the lowest dose examined was not present 8 days post-treatment, and that of the highest dose 15 days post-treatment, suggesting a gradual decline in effectiveness over time in females. Recent studies have demonstrated that single doses of psychedelics can produce long-lasting antidepressant effects in rodents subjected to the FST, primarily evaluated in adults. For example, a single dose of psilocybin improved FST performance in stressed male mice 15 days post-treatment (Sekssaoui et al., 2024). Sustained antidepressant-like effects were also observed in naïve adult male mice following a single injection of different psilocybin doses, with results lasting up to 7 days post-treatment (Zhao et al., 2024). Similarly, 5-MeO-DMT produced antidepressant-like effects 9 days post-treatment in adult male and female rats (Cameron et al., 2023), while psilocybin and LSD produced antidepressant-like responses lasting up to 5 weeks in a depression model using Wistar-Kyoto rats (Hibicke et al., 2020). Additional studies using selective bred rat strains as an intrinsic model of depression also reported long-term antidepressant-like properties after a single administration of psilocybin in adult male Wistar Han and Wistar-Kyoto rats (Kolasa et al., 2024). However, psilocybin and psilocin failed to induce antidepressant-like effects in the Flinders Sensitive Line rat model of depression, highlighting the need for alternative animal models and behavioural paradigms to fully evaluate psilocybin’s therapeutic potential (Jefsen et al., 2019).

While most studies examined single doses, repeated doses also showed promise. Intermittent chronic administration of DMT produced a strong antidepressant-like effect in adult male and female rats (Cameron and Olson, 2018). Similarly, repeated microdoses of psilocybin produced a sustained antidepressant effect 15 days post-administration in male and female mice subjected to a chronic stress (Sekssaoui et al., 2024). Another study demonstrated that repeated microdoses of psilocybin increased resistance to stress-induced anhedonia in adult male rats (Kiilerich et al., 2023). The novelty of our research has focused on characterizing, for the first time, the fast-acting antidepressant-like effects of psylocibin administered repeatedly during adolescence, revealing that both age and sex significantly influence treatment outcomes. In line with this, adolescence and female sex were associated with reduced antidepressant efficacy as observed with treatments such as ketamine, cannabidiol, fluoxetine, nortriptyline and electroconvulsive seizures in adolescent rats (Bis-Humbert et al., 2020; Bis-Humbert et al., 2021; García-Cabrerizo et al., 2020; Ledesma-Corvi et al., 2022; Ledesma-Corvi and García-Fuster, 2023). While these treatments improved certain affective-like behaviours in male adolescent rats, they were often ineffective in females and, in some cases, even detrimental during adolescence (García-Cabrerizo et al., 2020; Ledesma-Corvi et al., 2022). These findings emphasize the need to identify more effective treatments for adolescence by adapting paradigms to achieve better antidepressant efficacy, and the importance of considering sex differences, potentially influenced by the regulatory role of sex hormones (Ledesma-Corvi et al., 2023; Ledesma-Corvi and García-Fuster, 2023). In this context, our novel data proposes oral psylocibin to be further explored as a therapeutic option in sex-tailored clinical trials for adolescent patients with depression.

In conclusion, the present study demonstrates the rapid and long-lasting antidepressant efficacy of psilocybin in adolescent male and female rats. Overall, an acute oral administration was enough to show an antidepressant effect in both sexes at the doses used. However, sex differences were observed in the effectiveness of the antidepressant after repeated administrations, with the effects being more sustained in males for both tested doses as compared to females. Future research should investigate the mechanisms underlying the observed responses as well as the diminished antidepressant effectiveness in adolescent females after repeated treatment, while also evaluating safety. Additionally, exploring other doses or administration regimens that might enhance long-term antidepressant responses in females will be important. Regardless, psilocybin shows potential as a promising antidepressant with fast-acting and long-lasting effects during adolescence, a stage where the effectiveness of conventional antidepressants and available pharmacological options is often uncertain. Notably, the results from oral administration provide valuable translational insights, as this common route of psilocybin consumption accounts for important pharmacokinetic factors related to its potential antidepressant effects.

